# Phenolic degradation by catechol dioxygenases is associated with pathogenic fungi with a necrotrophic lifestyle in the Ceratocystidaceae

**DOI:** 10.1101/2021.12.30.474597

**Authors:** Nicole C Soal, Martin HA Coetzee, Magriet A van der Nest, Almuth Hammerbacher, Brenda D Wingfield

## Abstract

Fungal species of the Ceratocystidaceae grow on their host plants using a variety of different lifestyles, from saprophytic to highly pathogenic. Although many genomes of fungi in the Ceratocystidaceae are publicly available, it is not known how the genes that encode catechol dioxygenases (*CDOs),* enzymes involved in the degradation of phenolic plant defence compounds, differ among members of the Ceratocystidaceae. The aim of this study was therefore to identify and characterize the genes encoding CDOs in the genomes of Ceratocystidaceae representatives. We found that genes encoding CDOs are more abundant in pathogenic necrotrophic species of the Ceratocystidaceae and less abundant in saprophytic species. The loss of the *CDO* genes and the associated 3-oxoadipate catabolic pathway appears to have occurred in a lineage-specific manner. Taken together, this study revealed a positive association between *CDO* gene copy number and fungal lifestyle in Ceratocystidaceae representatives.

## Introduction

The interaction between plants and pathogenic fungi is complex, with the continuous coevolution of plant defence mechanisms and virulence mechanisms of pathogens (Anderson et al. 2010). Upon invasion by a pathogen, plants defend themselves from infection through various mechanisms, such as the formation of physical barriers, synthesis of antimicrobial proteins and chemical defence compounds (Jones and Dangl 2006; Ferreira et al. 2007). Plants synthesise a variety of chemical compounds (terpenes, alkaloids and different types of phenolics) that are either preformed or produced upon infection (Lattanzio et al. 2006; Lattanzio et al. 2008). Some of these phenolic compounds include known antifungal agents, such as catechin, stilbenes, isoflavonoids and condensed tannins (Jeandet et al. 2002; Kocaçalişkan et al. 2006; Lattanzio et al. 2006; Liu et al. 2017; Ullah et al. 2017).

Phytopathogenic fungi have evolved to overcome host defence responses. For example, some fungi can degrade phenolics that are part of the host defence response via specialised enzymatic pathways (Westrick et al. 2021). In many instances the phenolic degradation pathways in fungi are clustered and connected by key classes of enzymes (Gluck-Thaler et al. 2018; Gluck-Thaler and Slot 2018). This includes the enzyme class catechol dioxygenases (CDO), which can cleave aromatic rings at the catecholic bond through the addition of molecular oxygen (Hayaishi 1966; Broderick 1999). The CDO enzymes are divided into two families either catalysing the first reaction of the *ortho*-cleavage or *meta*-cleavage pathways. These enzyme families have evolved independently from one another and are found in both eukaryotic and prokaryotic microorganisms (Harayama and Rekik 1989; Eltis and Bolin 1996; Vaillancourt et al. 2006).

The intradiol dioxygenases (EC 1.13.11.1), are part of the catechol branch of the 3-oxoadiptae pathway (also known as the *ortho-*cleavage pathway or the β-ketoadipate pathway) (Harwood and Parales 1996; Martins et al. 2015). Intradiol dioxygenases cleave the carbon-carbon bond of catecholic aromatic compounds between the two adjacent hydroxy substituents (Figure 1) (Broderick 1999; Vaillancourt et al. 2006). In this pathway, the catecholic substrate is cleaved to form a muconolactone (Harwood and Parales 1996), which is further broken down via multiple enzymatic reactions to form the products succinyl-CoA and acetyl-CoA which enter the tricarboxylic acid cycle and provide substrates for the production of the high-energy molecules, NADH and ATP (Figure 1) (Ornston and Stanier 1966; Harwood and Parales 1996; Martins et al. 2015). In the *meta-*cleavage pathway, on the other hand, an extradiol dioxygenase cleaves the catecholic substrate adjacent to a hydroxy substituent to form a muconate semialdehyde (Vaillancourt et al. 2006). This compound is further broken down to form pyruvate and acetaldehyde which are metabolised by the glycolysis pathway (Figure 1) (Maruyama et al. 2004).

**Figure 1:**
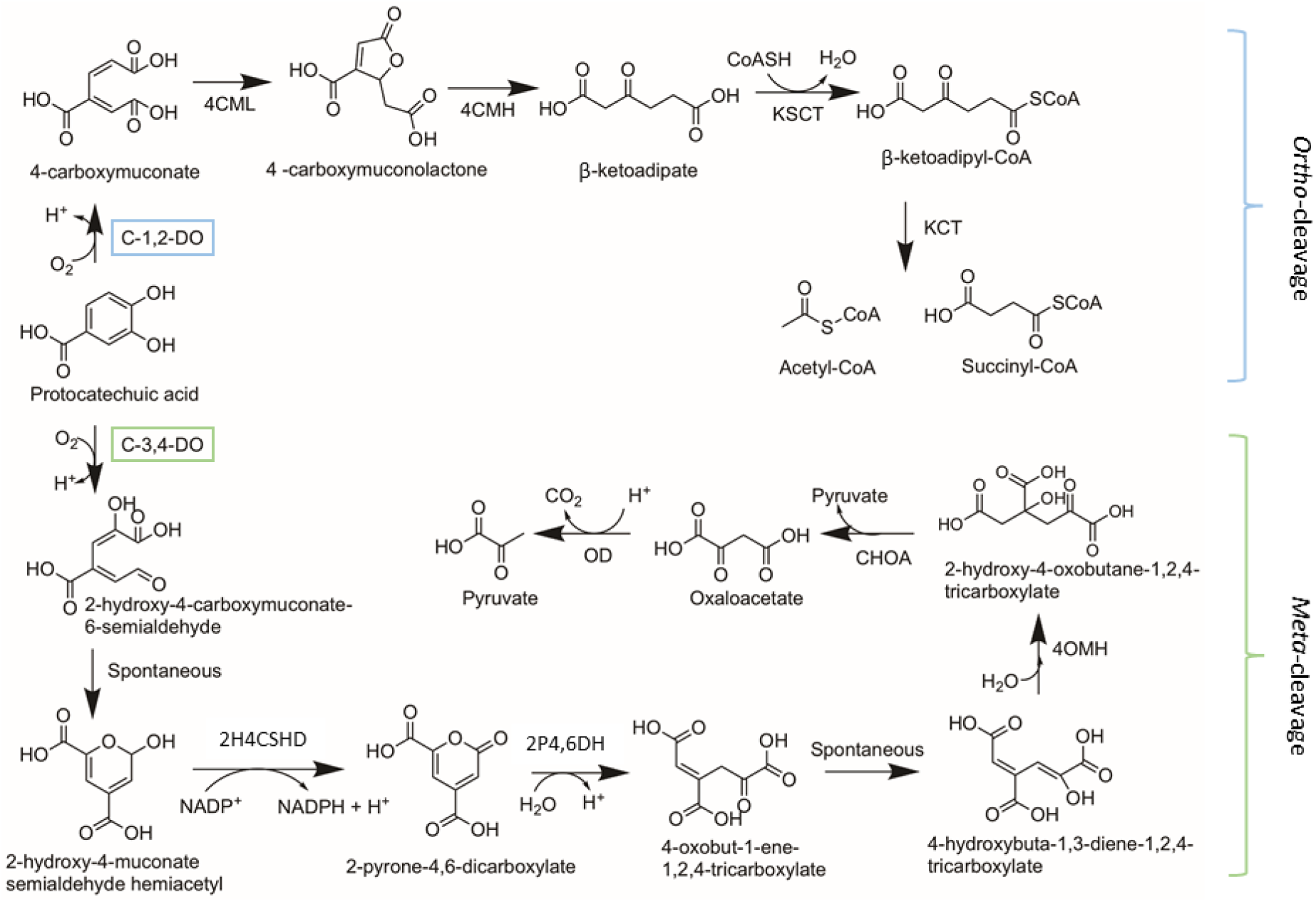
The *ortho-* and *meta-* cleavage routes of phenolic degradation by microbial species. Blue bracket over C-1,2-DO represents the step in the *ortho-*cleavage pathway that CDO1, 2 and 4 are involved in. Green bracket over C-3,4-DO indicated the step in the *meta*-cleavage pathway that CDO3 is involved in. (Enzymes: C-1,2- DO – catechol 1,2-dioxygenase, 4CML – 4-carboxymuconate lactonase, 4CMH – 4- carboxymuconolactone hydrolase, KSCT – β-ketoadipate:succinyl-CoA transferase, KCT – β-ketoadipate-CoA thiolase; C-3,4-DO – catechol 3,4-dioxygenase, 2H4CSHD – 2-hydroxy-4-carboxymuconate-6-semialdehyde dehydrogenase, 2P4,6DH – 2-pyrone-4,6-dicarboxylate hydrolase, 4OMH – 4-oxalomesaconate hydratase, CHOA – 4-carboxy-4-hydroxy-2-oxoadipate aldolase, OD – oxaloacetate β-decarboxylase). Chemical structures and pathways, adapted from Wadke *et al*. (2016).

Fungi are thus able to utilise phenolic plant defence metabolites as carbon sources through the use of CDOs and their respective catabolic pathways (Camarero et al. 1994; Bugg et al. 2011). The action of CDOs during plant infection has been shown to contribute to the pathogenicity and virulence of some phytopathogenic fungi (Shanmugam et al. 2010; Hammerbacher et al. 2013; Wadke et al. 2016). For example, the fungal maize pathogen, *Cochliobolus heterostrophus*, produces an intradiol CDO in response to phenolic compounds present in its environment (Shanmugam et al. 2010). Members of the family Ceratocystidaceae utilize CDOs to overcome host responses. For example, the pathogen *Endoconidiophora polonica* makes use of CDOs in the degradation of flavonoids and stilbenes produced by its conifer host (Wadke et al. 2016). Previous studies revealed that increased expression of the *CDO* genes in this fungus increased virulence of the pathogen (Hammerbacher et al. 2013; Wadke et al. 2016).

In spite of the importance of CDOs in the plant host-fungal pathogen interaction, very little is known about CDOs in the Ceratocystidaceae. This fungal family include agricultural crop pathogens such as the sweet potato black rot pathogen, *Ceratocystis fimbriata,* the banana crown rot pathogen, *Thielaviopsis musarum,* and the causal agent of carrot root rot, *Berkeleyomyces basicola* (Halsted and Fairchild 1891; Melo et al. 2016; Nel et al. 2017). There are also many species which cause disease on important forest trees, including the conifer blue-stain fungi, *Endoconidiophora polonica* and *E. laricicola,* the wattle wilt pathogen, *Ce. albifundus,* and the causal agent of sapstreak in sugar maple trees, *Davidsoniella virescens* (Krokene and Solheim 1998; Roux et al. 2007; Richter 2012). Apart from these pathogenic species there are also a number of saprophytic species in the Ceratocystidaceae that do not cause disease or which are considered weakly pathogenic. These include *Huntiella* species, particularly *H. moniliformis* and *H. bhutanensis* (Wilson et al. 2015; Wingfield, Duong, et al. 2016), weakly pathogenic fungi such as *D. eucalypti,* found on *Eucalyptus* (Kile et al. 1996), and the ambrosia beetle symbionts, *Ambrosiella xylebori* and *A. cleistominuta* (von Arx and Hennebert 1965; Mayers et al. 2019).

There are a large number of sequenced genomes in the Ceratocystidaceae (Wilken et al. 2013; Van der Nest et al. 2014; Wingfield et al. 2015; Wingfield, Ambler, et al. 2016; Wingfield et al. 2017) which provide an important resource for determining the presence of these enzymes across different genera. However, to date, only four CDOs were identified in *E. polonica* (Wadke et al. 2016). It is still unknown whether other members of the Ceratocystidaceae with different ecological lifestyles have similar enzymes. The aim of this study was to identify and characterize the *CDOs* in pathogenic and non-pathogenic species of the Ceratocystidaceae, as well as understand the evolutionary history of the catechol dioxygenases and their involvement in fungal nutrition.

## Materials and methods

### Identification and characterisation of putative CDOs in the Ceratocystidaceae

To identify putative CDO homologs present in the genomes of the fungal species, we used the sequences identified in *E. polonica* to search the publicly available genomes. These included the nucleotide sequences of three intradiol dioxygenases (*EpCDO1*, GenBank Accession: KU221039; *EpCDO2*, GenBank Accession: KU221040; *EpCDO4*, GenBank Accession: KU221042) and one extradiol dioxygenase (*EpCDO3*, GenBank Accession: KU221041) (Wadke et al. 2016).

Putative CDO homologs were identified using local BLAST searches (tBLASTn, expect (E)-values > 10^5^) using CLC Genomics Workbench v 8.1 (https://www.qiagenbioinformatics.com/). The corresponding scaffold or contig on which there was a hit was extracted in CLC Main Workbench and open reading frames were predicted using Web AUGUSTUS (http://bioinf.unigreifswald.de/augustus/) (Hoff et al. 2012; Hoff and Stanke 2013). The genes identified from the annotations were confirmed with a reciprocal BLASTp on NCBI. The functional domains of the CDOs were annotated using InterProScan (Jones et al. 2014), the conserved domains were identified using the Conserved Domain (CD) function on NCBI (http://www.ncbi.nlm.-nih.gov/Structure/cdd/wrpsb.cgi). The predicted proteins were also assessed for transmembrane domains using TMHMM (Krogh et al. 2001) and for signal peptides using SignalP (Petersen et al. 2011). Orthologous proteins were identified using OrthoMCL (Li et al. 2003).

### Phylogenetic analysis

A phylogenetic approach was taken to determine if the genes could be considered orthologous and to determine the evolutionary history and relationships of the respective genes. The genome sequence information for 30 species of the Ceratocystidaceae was used, in addition the genome sequences from *Fusarium circinatum* and *F. fujikuroi* were used to provide outgroup gene sequences (Table 1). They were obtained from the GenBank database of the National Centre for Biotechnology Information (NCBI; https://www.ncbi.nlm.nih.gov/).

**Table 1:**
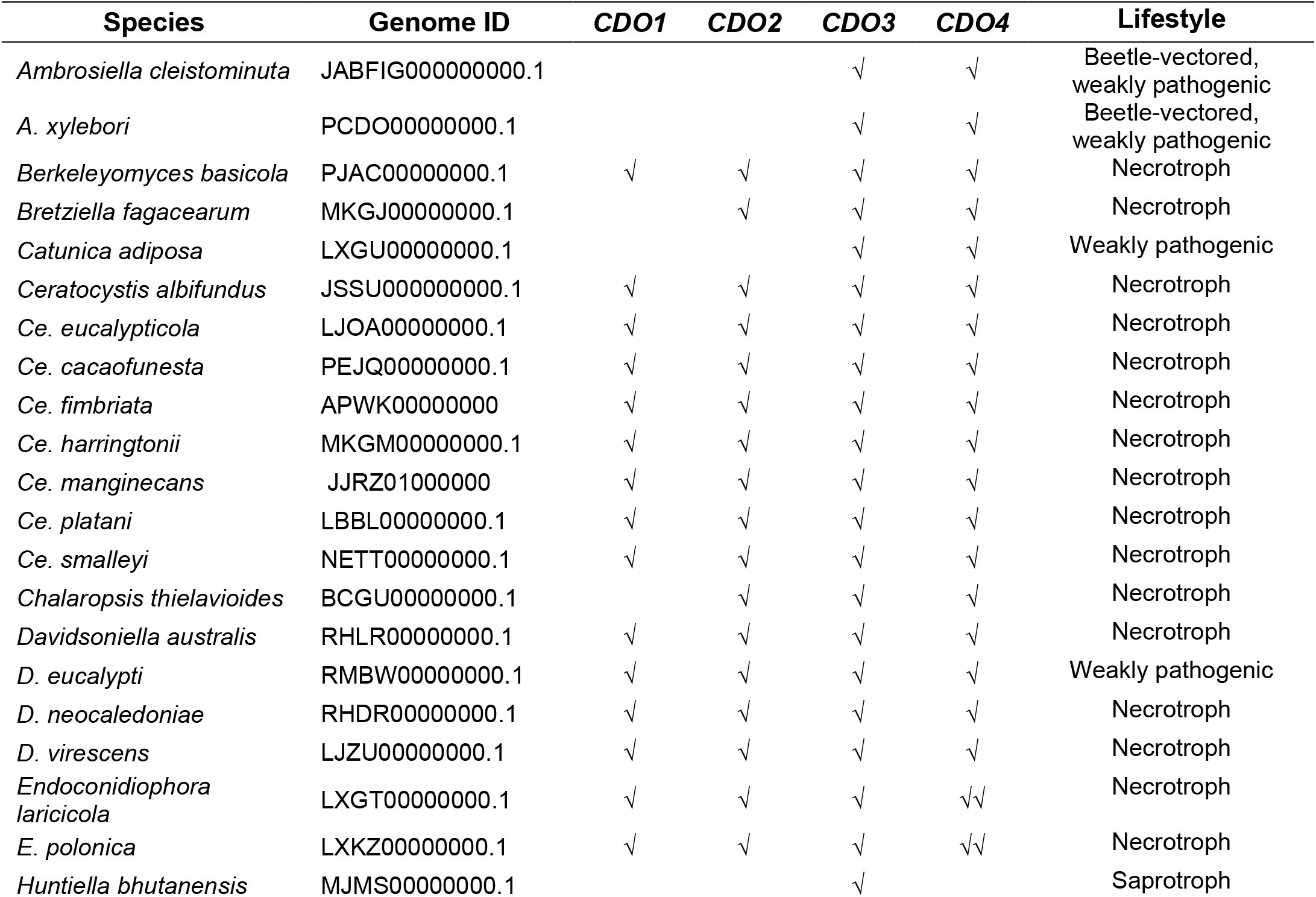
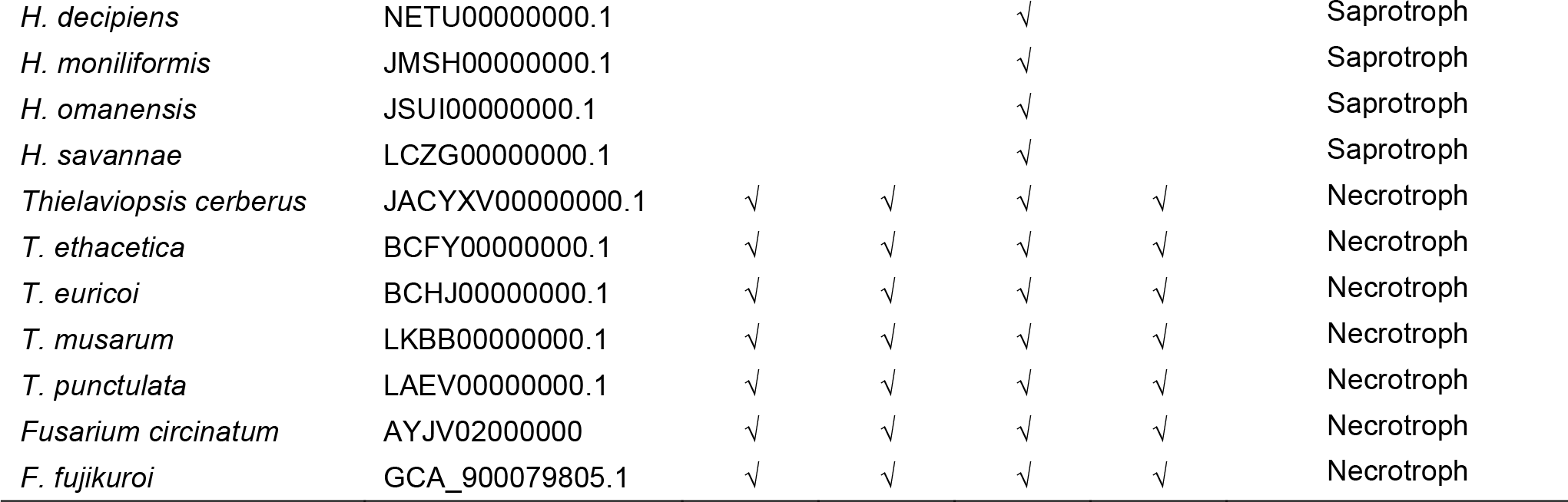
Presence of the CDO 1-4 gene transcripts across the Ceratocystidaceae with the genome information and ecological lifestyle of each species used in this

| Species | Genome ID | *CDO1* | *CDO2* | *CDO3* | *CDO4* | Lifestyle |
| --- | --- | --- | --- | --- | --- | --- |
| *Ambrosiella cleistominuta* | JABFIG000000000.1 |  |  | √ | √ | Beetle-vectored,<br>weakly pathogenic |
| *A. xylebori* | PCDO000000000.1 |  |  | √ | √ | Beetle-vectored,<br>weakly pathogenic |
| *Berkeleyomyces basicola* | PJAC000000000.1 | √ | √ | √ | √ | Necrotroph |
| *Bretziella fagacearum* | MKGJ000000000.1 |  | √ | √ | √ | Necrotroph |
| *Catunica adiposa* | LXGU000000000.1 |  |  | √ | √ | Weakly pathogenic |
| *Ceratocystis albifundus* | JSSU000000000.1 | √ | √ | √ | √ | Necrotroph |
| *Ce. eucalypticola* | LJOA000000000.1 | √ | √ | √ | √ | Necrotroph |
| *Ce. cacaofunesta* | PEJQ000000000.1 | √ | √ | √ | √ | Necrotroph |
| *Ce. fimbriata* | APWK000000000 | √ | √ | √ | √ | Necrotroph |
| *Ce. harringtonii* | MKGM000000000.1 | √ | √ | √ | √ | Necrotroph |
| *Ce. manginecans* | JJRZ01000000 | √ | √ | √ | √ | Necrotroph |
| *Ce. platani* | LBBL000000000.1 | √ | √ | √ | √ | Necrotroph |
| *Ce. smalleyi* | NETT000000000.1 | √ | √ | √ | √ | Necrotroph |
| *Chalaropsis thielavioides* | BCGU000000000.1 |  | √ | √ | √ | Necrotroph |
| *Davidsoniella australis* | RHLR000000000.1 | √ | √ | √ | √ | Necrotroph |
| *D. eucalypti* | RMBW000000000.1 | √ | √ | √ | √ | Weakly pathogenic |
| *D. neocaledoniae* | RHDR000000000.1 | √ | √ | √ | √ | Necrotroph |
| *D. virescens* | LJZU000000000.1 | √ | √ | √ | √ | Necrotroph |
| *Endoconidiophora laricicola* | LXGT000000000.1 | √ | √ | √ | √√ | Necrotroph |
| *E. polonica* | LXKZ000000000.1 | √ | √ | √ | √√ | Necrotroph |
| *Huntiella bhutanensis* | MJMS000000000.1 |  |  | √ |  | Saprotroph |
| *H. decipiens* | NETU00000000.1 |  |  | √ |  | Saprotroph |
| *H. moniliformis* | JMSH00000000.1 |  |  | √ |  | Saprotroph |
| *H. omanensis* | JSUI00000000.1 |  |  | √ |  | Saprotroph |
| *H. savannae* | LCZG00000000.1 |  |  | √ |  | Saprotroph |
| *Thielaviopsis cerberus* | JACYXV00000000.1 | √ | √ | √ | √ | Necrotroph |
| *T. ethacetica* | BCFY00000000.1 | √ | √ | √ | √ | Necrotroph |
| *T. euricoi* | BCHJ00000000.1 | √ | √ | √ | √ | Necrotroph |
| *T. musarum* | LKBB00000000.1 | √ | √ | √ | √ | Necrotroph |
| *T. punctulata* | LAEV00000000.1 | √ | √ | √ | √ | Necrotroph |
| *Fusarium circinatum* | AYJV02000000 | √ | √ | √ | √ | Necrotroph |
| *F. fujikuroi* | GCA_900079805.1 | √ | √ | √ | √ | Necrotroph |

The amino acid sequences for each of the individual genes (CDO1-4) were aligned using MAFFT (Multiple sequence alignment based on fast Fourier transform) v 7 (https://mafft.cbrc.jp/alignment/server/) using the L-INS-i option (Katoh and Toh 2008) to generate a single alignment per gene. The alignments were trimmed and in the case of CDO4, the N-terminal signal peptide was excluded. The alignments were then analysed using a Maximum Likelihood (ML) method with RAxML v8 (Stamatakis 2014). The predicted best fit substitution model for amino acid data as indicated by ProtTest 3 (Darriba et al. 2011) was incorporated in the ML analyses. The evolutionary models were the Whelan and Goldman (Whelan and Goldman 2001) with gamma distribution (WAG+G) for CDO1 and CDO2 data, and the Le-Gascuel model (Le and Gascuel 2008) with gamma distribution (LG+G) for CDO3 and CDO4. The branch support was determined using 1000 bootstrap replicates. The phylogenetic trees were viewed using MEGA v.7 (Kumar et al. 2018) and FigTree v1.3.1 (http://tree.bio.ed.ac.uk/software/figtree/).

A multi locus species tree was generated for all the species used in this study. The nucleotide sequences for the Minichromosome Maintenance Complex Component 7 (MCM7), DNA-directed RNA polymerase II subunit RPB1 and RPB2 (RPB1, RPB2) and Elongation Factor 2 (EF2) genes were identified in the genomes using the BLASTn function in CLC Main Workbench v 8.1. The nucleotide sequences were aligned using MAFFT v 7 and concatenated using FASconCAT-G v 1.04 (Kück and Longo 2014). The alignments were converted to Phylip files using PAUP v 4.0a (Swofford 2002). An ML analysis was conducted on the data using the predicted evolutionary model of a General Time Reversible with gamma distribution (GTR+G) as determined with jModelTest (Darriba et al. 2012). RAxML v8 was used to perform the ML analysis, with the data being partitioned into the respective loci, and the parameters for each partition allowed to vary independently. Bootstrap support was determined with 1000 rapid bootstrap replicates. The tree was viewed using viewed using MEGA v.7 and FigTree v1.3.1.

### Gene synteny flanking *CDOs*

The gene synteny surrounding the *CDO* genes was investigated to determine whether the *CDOs* are found in specialised enzymatic clusters. For each of the *CDOs*, the five genes flanking either side of the genes, where available, were identified using BLASTx searches on NCBI. These flanking regions were assessed for synteny using EasyFIG 2.2.3 (Sullivan et al. 2011). This was done to identify any possible gene clusters involved in the degradation of catechol derivatives, and to determine whether there were regions in the surrounding genes that were indicative of gene expansions, gains or losses. The programme uses a BLASTn function to compare the sequences of the input fasta files using a similarity threshold with a maximum E value of 0.001.

**Figure 2:**
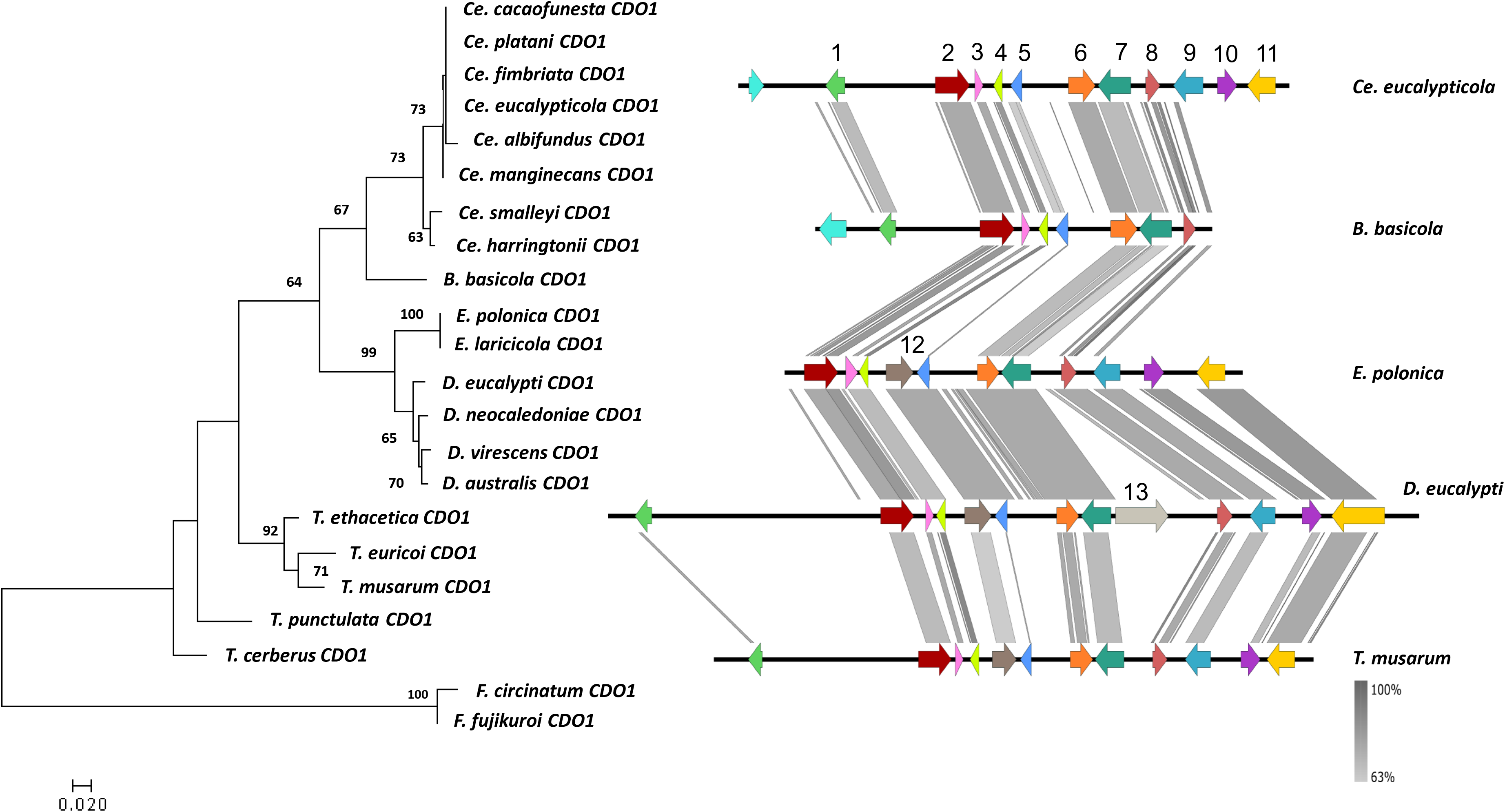
Maximum likelihood gene tree (left) for CDO1 amino acid sequences (1000 bootstrap replicates) with outgroup species *F. circinatum* and *F. fujikuroi*. *S*ynteny map (right) for representative species of each of the genera present in the gene tree (EasyFig) representing the sequence similarity of the genomic regions flanking *CDO1*. \**CDO1* (6, orange) including five genes upstream and downstream of the *CDO*. Genes which are numbered and coloured are shown to be syntenic and conserved through the majority of the species, with some unique genes also included. The genes without numbers or coloured grey were not found to be syntenic across genera. From the left: 1 – hypothetical protein; 2 – putative phosphatidylinositol N-acetylglucosaminyltransferase subunit C; 3 – mediator of RNA polymerase II transcription subunit 31; 4 – DNA-directed RNA polymerase III subunit RPC10; 5 – hypothetical protein containing CFEM domain; 6 – *CDO1*; 7 – tRNA (uracil-O(2)-)-methyltransferase; 8 – eukaryotic translation initiation factor eIF-1; 9 – N-glycosylation protein; 10 – hypothetical protein; 11 – sodium/pantothenate symporter; 12 – Ser/Thr protein phosphatase family protein; 13 – reverse transcriptase.

### Analysis of *CDO* gene family evolution

The gene family contractions and expansions across the Ceratocystidaceae were inferred using the Count software package (http://www.iro.umontreal.ca/~csuros/gene_content/count.html) through inferring the character’s history given the phylogeny (Csűrös and Miklós 2006; Csűrös 2010). The analysis was performed using a Wagner parsimony method that penalises individual family members for their loss (loss penalty = 1) and gains, inferring the history with a minimum penalty on the previously described multi locus species tree generated. To further analyse the data, the ancestral character states were constructed for each individual gene using Mesquite v 3.0.4 (Maddison and Maddison 2019). The analyses were based on Parsimony reconstruction and used the tree generated from the combined data (see above) as the phylogenetic hypothesis.

### Identification of the *ortho-* and *meta-*cleavage pathways in the Ceratocystidaceae

To identify the putative homologs of the *ortho-* and *meta-*cleavage pathways, the characterised genes of the pathways from closely related species were used in a tBLASTn search against the 30 genomes of the Ceratocystidaceae (Table 1) using CLC Genomics Workbench v 8.1. The reference genes used as queries include carboxy-cis,cis-muconate cyclase (Ce. fimbriata PHH5432.1); 3-oxoadipate enol-lactonase (Ce. platani KKF95260.1); 3-oxoadipate CoA-transferase (Ce. platani KKF92536.1); acetyl-CoA acyltransferase (Ce. fimbriata PHH52801.1). The corresponding scaffold or contig on which a gene was identified was extracted in CLC Main Workbench and open reading frames were predicted using web AUGUSTUS. The genes identified from the annotations were confirmed with a reciprocal BLASTp on NCBI.

### Statistical analysis

The data was separated into pathogens and non-pathogens (as seen in Table 1), and the number of species in each category were further assigned to a category of i) having all four *CDOs* present or ii) not having all four *CDOs* present. This count data was subjected to a Fisher’s Exact Test using a 95% confidence interval (two-tailed), with an alternate hypothesis that the true odds ratio is not equal to 1. All analyses were conducted using R (www.r-project.org).

## Results

### Identification and characterisation of putative CDOs in the Ceratocystidaceae

From the 32 genomes used in this study (Table 1), putative CDOs (*CDO1* – *CDO4*) were identified based on their sequence identity to previously identified genes from *E. polonica* (Wadke et al. 2016). The genes identified were consistent between species within the respective genera (Table 1). Only one of the genes, *CDO3*, was identified in all the species included in this study (Table 1). *Ceratocystis* spp., *Davidsoniella* spp., one *Berkeleyomyces* sp. and *Thielaviopsis* spp. had single copies of all four of the previously described *CDO* genes. *Bretziella fagacearum* and *Chalaropsis thielavioides* genomes contained gene sequences for *CDO2, 3* and *4*. *Ambrosiella* spp. and *Ca. adiposa* contained gene sequences for *CDO3* and *CDO4*. Species of *Huntiella* had only one extradiol dioxygenase, *CDO3*. We also identified a gene duplication of *CDO4* in *E. polonica* and *E. laricicola*. The genomes of the outgroup species *F. circinatum* and *F. fujikuroi* contained one copy of each of the CDOs (Table 1).

The genome architecture of *CDO1* using InterProScan, CD and SignalP identified a conserved intradiol ring-cleavage dioxygenase core domain (IPR015889) adjacent to a conserved intradiol ring-cleavage dioxygenase C-terminal (IPR000627). The sequences contained the conserved residues of the active site and dimer interface, including the residues involved in binding the iron(III) cofactor.

The predicted conserved domains identified in the *CDO2* genes included an N-terminal domain (IPR007535) characteristic of an intradiol cleaving dioxygenase and an intradiol ring-cleavage dioxygenase core domain (IPR015889). A third intradiol dioxygenase was identified, *CDO4*, was identified. All the *CDO4* homologs contained an N-terminal signal peptide as predicted by InterProScan and SignalP, which was consistent with findings by Wadke *et al*. (2016). They were well conserved and spanned 20 amino acids into the gene. Thus, the CDO4 protein is predicted to be secreted to the extracellular space. The analyses identified the ASA-HP cleavage site between amino acids 20 and 21. This is a standard secretory peptide transported by the Sec translocon protein and cleaved by Signal Peptidase I (Auclair et al. 2012). This was consistent in each *CDO4* gene included in this study. The intradiol ring-cleaving dioxygenase core C-terminal domain (IPR000627) was present in the each of the *CDO4* transcripts.

One extradiol cleaving dioxygenase, *CDO3*, was identified. This extradiol ring-cleavage dioxygenase was similar to a class III enzyme, with a conserved subunit B domain (IPR004183), with similarity to a 4,5-DOPA extradiol dioxygenase (EC 1.13.11.29). A gene duplication of *CDO4* was identified in the *E. polonica* and *E. laricicola* genomes. The genes were located on different scaffolds, and were surrounded by different flanking genes. The sequence similarity of the coding regions of the two genes was 92.73% in *E. polonica* and 93.9% in *E. laricicola* (Figure S1). When the amino acid sequences of the homologous *CDO4* genes at the same genomic location were compared to one another, they were 99.13% similar in percentage identity to each other (Figure S1). The duplicate gene is located on scaffolds that are 24 364 bp and 24 420 bp in length in *E. polonica* and *E. laricicola*, respectively (Wingfield, Ambler, et al. 2016). Each scaffold had three genes with a predicted ORF present. The other genes present on the scaffold are predicted to be an isoamyl alcohol oxidase/dehydrogenase and a hypothetical protein. The cleavage site for the secretory signal was identified in the *Endoconidiophora CDO4* gene duplicates.

### Phylogenetic analysis

The ML phylogenies of the individual genes showed close congruency to the species tree, and each gene formed a clade that included members of the same genus (Figures 2-5). The duplicated *CDO4* genes identified in the *Endoconidiophora* species grouped with the *CDO4* genes identified previously (Figure 5), but formed their own clade, suggesting that the gene expansion occurred in an ancestral *Endoconidiophora* lineage.

### Gene synteny flanking *CDOs*

The genes flanking the *CDOs* differed between genera. Within-genus comparisons revealed high conservation of gene order and orientation. When compared across genera, there were certain genera that had conserved gene order and orientation, such as *Ceratocystis* and *Davidsoniella* (Figures 2-5). However, there were some outliers, such as *Br. fagacearum* and the genus *Huntiella*, showing low synteny compared to the other genera included in the dataset (Figures 2-5).

The gene orientation and order for *CDO1* showed gene synteny between the *Ceratocystis* species and the *Berkeleyomyces* species (Figure 2). There are many similarities between the genera, with only a few genes that differed between them. A similar pattern is observed for *CDO2*, with genes downstream of the *CDO2* gene showing high synteny between the genera, whereas, upstream of the gene, the gene order is less conserved in *Thielaviopsis* (Figure 3). One of the genes upstream of *CDO2* included a carboxy-*cis,cis-*muconate cyclase (see gene 5 in Figure 3) which is one of the enzymes used in the catechol branch of the *ortho-*cleavage pathway. The genes upstream of *CDO3* were more conserved than those downstream in some of the species. There were striking differences between the flanking genes of *CDO3* of *Huntiella moniliformis* compared to the rest of the species in the other genera (Figure 4).

**Figure 3:**
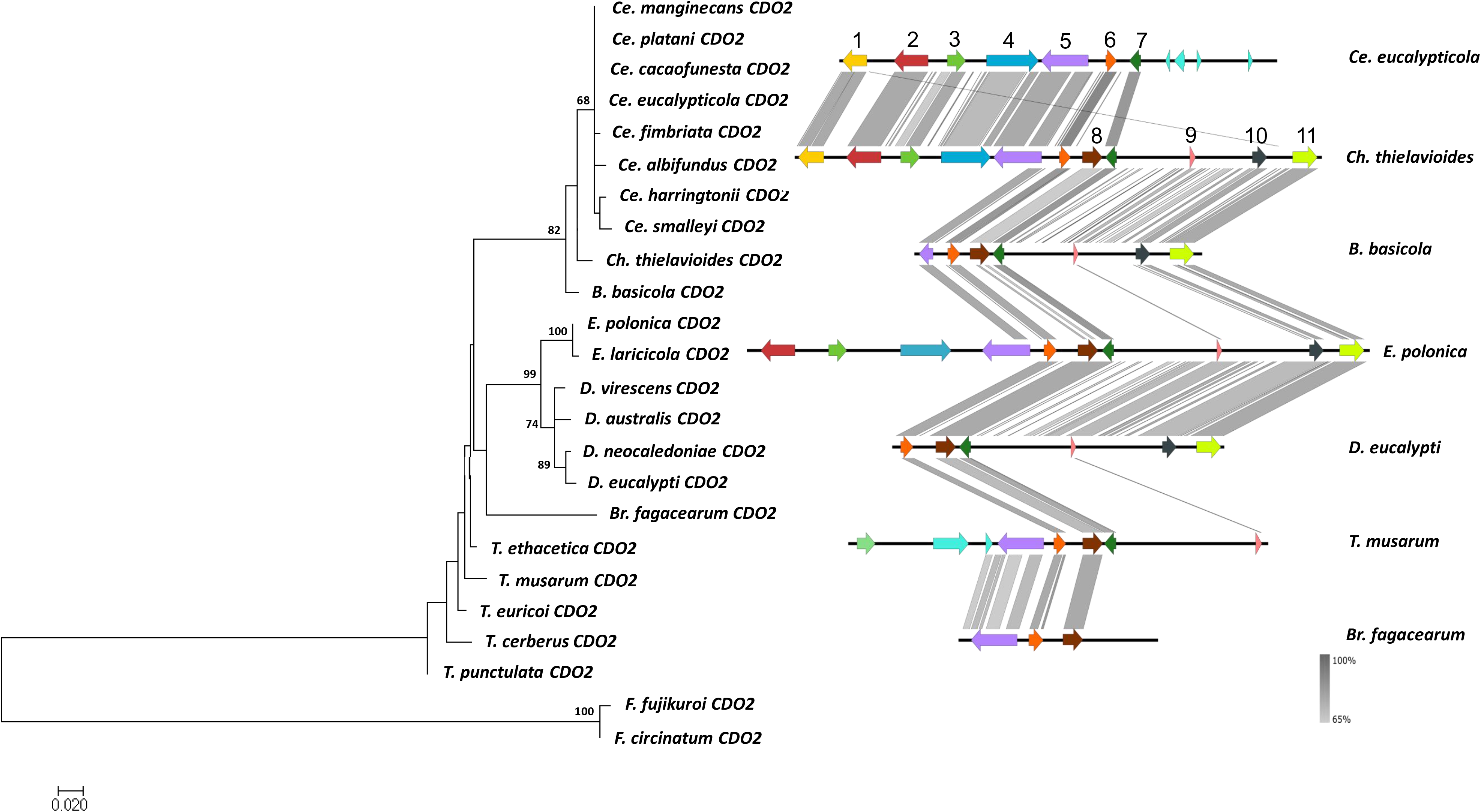
Maximum likelihood gene tree (left) for CDO2 amino acid sequences (1000 bootstrap replicates) with outgroup species *F. circinatum* and *F. fujikuroi. S*ynteny map (right) for representative species of each of the genera present in the gene tree (EasyFig) representing the sequence similarity of the genomic regions flanking *CDO2*. \**CDO2* (6, orange) including five genes upstream and downstream of the *CDO*. Genes which are numbered and coloured are shown to be syntenic and conserved through the majority of the species, with some unique genes also included. The genes without numbers or coloured grey were not found to be syntenic across genera. From the left: 1 – neutral ceramidase; 2 – ATP-dependent DNA helicase srs2; 3 – hypothetical protein containing DUF 4591 domain; 4 – sterol regulatory element-binding protein 1; 5 – carboxy-*cis,cis-*muconate cyclase; 6 – *CDO2*; 7 – nitronate monooxygenase; 8 – endonuclease/exonuclease/phosphatase, COG 2374; 9 – double-stranded RNA binding motif; 10 – 37S ribosomal protein S7 mitochondrial; 11 – hypothetical protein.

**Figure 4:**
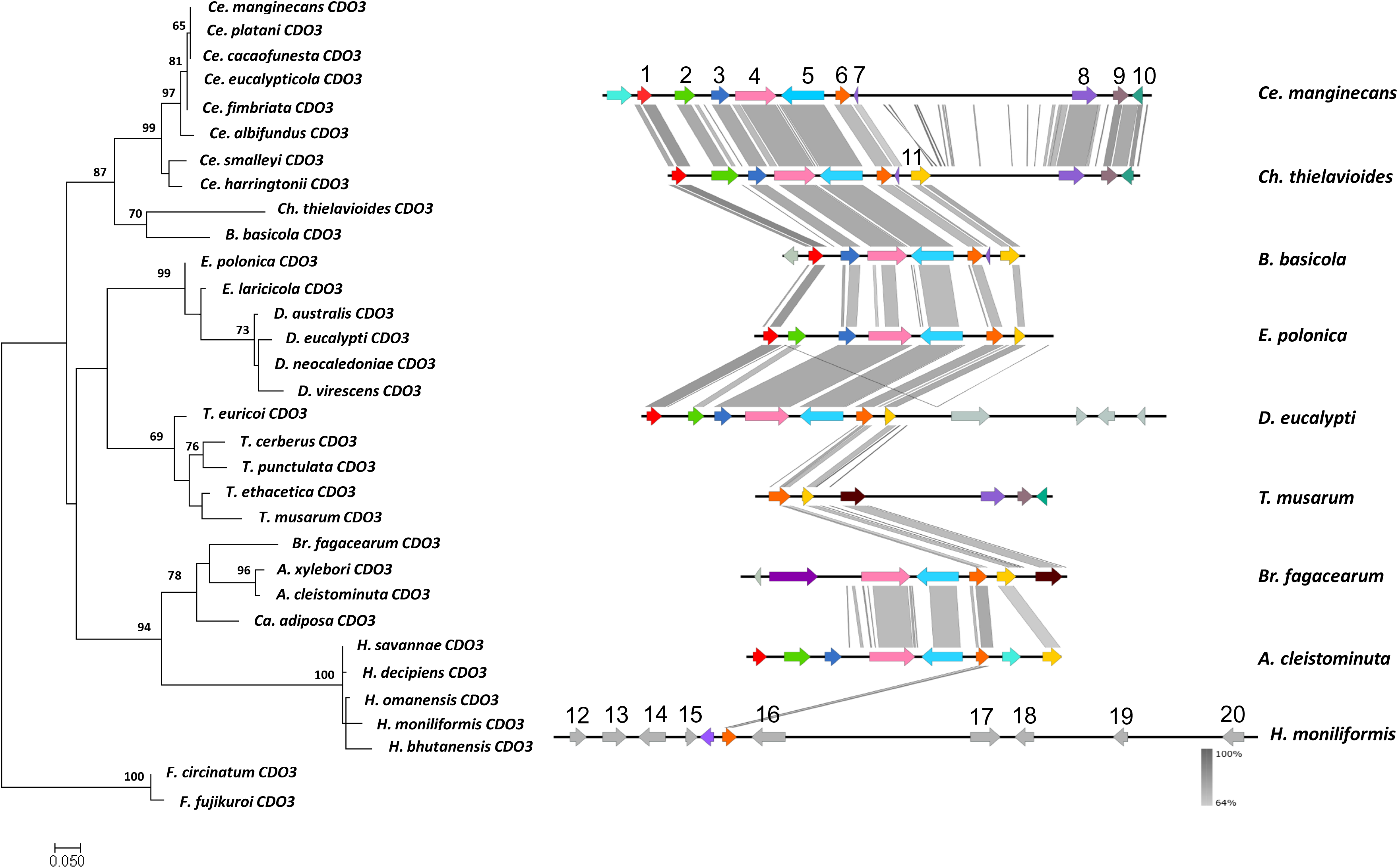
Maximum likelihood gene tree (left) for CDO3 amino acid sequences (1000 bootstrap replicates) with outgroup species *F. circinatum* and *F. fujikuroi*. *S*ynteny map (right) for representative species of each of the genera present in the gene tree (EasyFig) representing the sequence similarity of the genomic regions flanking *CDO3*. \**CDO3* (6, orange) including five genes upstream and downstream of the *CDO*. Genes which are numbered and coloured are shown to be syntenic and conserved through the majority of the species, with some unique genes also included. The genes without numbers or coloured grey were not found to be syntenic across genera. From the left: 1 – 40S ribosomal protein S19; 2 – hypothetical protein; 3 – fluoride export protein 1; 4 – hypothetical protein containing DUFF 4449 domain; 5 – DNA ligase 4; 6 – *CDO3*; 7 – hypothetical protein; 8 – COP9 signalosome complex subunit 5; 9 – hypothetical protein; 10 – putative RNA-binding protein sce3; 11 – hypothetical protein; 12 – universal stress protein A family protein C25B2.10; 13 – GPCR-type G protein 2; 14 – putative RNA-binding protein sce3; 15 – hypothetical protein; 16 – putative vacuolar membrane protein; 17 – glycerol kinase; 18 – mannose-1-phosphate guanyltransferase; 19 – hypothetical protein; 20 – protein farnesyltransferase subunit beta.

In many genera the genes flanking *CDO4* are predicted to be involved in the shikimate pathway and the catabolism of quinate, a product of the shikimate pathway (Figure 5). Quinate is degraded by various enzymes to a form that can enter the β- ketoadipate pathway. The genes identified as flanking *CDO4* include a catabolic 3- dehydroquinase (EC 1.1.5.8), 3-dehydroshikimate dehydratase (EC 4.2.1.10), quinate dehydrogenase (EC 1.1.1.282) and a quinate permease (Pf00083) which collectively are involved in the transformation of quinic acid to protocatechuate. This gene cluster was not present in *Br. fagacearum* and the *Ambrosiella* species (Figure 5). There were also large differences in the gene orientation and order between the different species for the genes flanking *CDO4* (Figure 5), as the syntenic genes found in the *Ambrosiella* species were dispersed in a different order, with some of the genes appearing upstream of *CDO4* on the contig. In *Br. fagacearum* and the genus *Ambrosiella* only one syntenic gene was found on the scaffold (Histone-lysine N-methyltransferase SET9), where the rest of the genes lacked synteny to the other genera (Figure 5).

**Figure 5:**
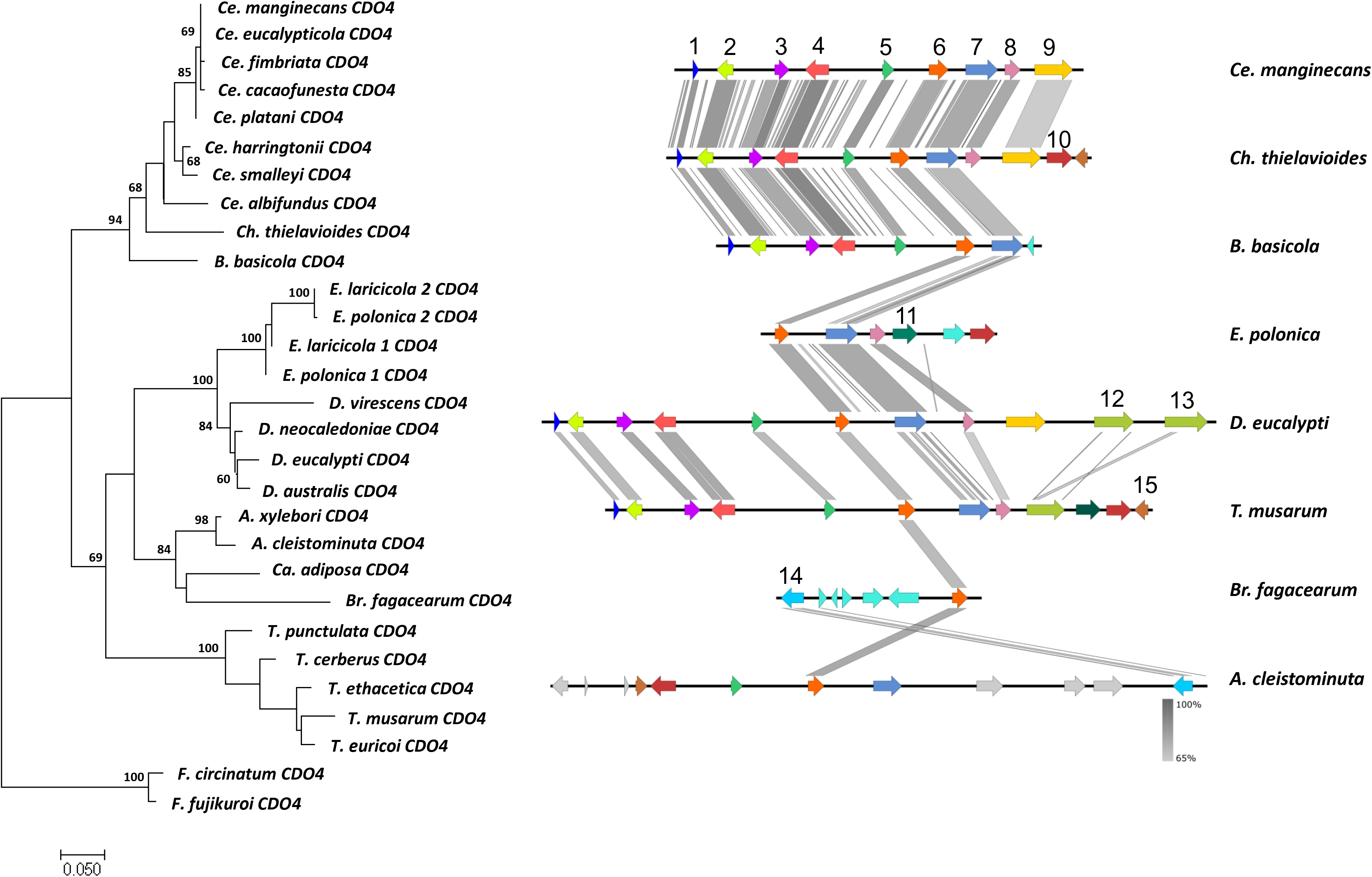
Maximum likelihood gene tree (left) for CDO4 amino acid sequences (1000 bootstrap replicates) with outgroup species *F. circinatum* and *F. fujikuroi. S*ynteny map (right) for representative species of each of the genera present in the gene tree (EasyFig) representing the sequence similarity of the genomic regions flanking *CDO4*. \**CDO4* (6, orange) including five genes upstream and downstream of the *CDO*. Genes which are numbered and coloured are shown to be syntenic and conserved through the majority of the species, with some unique genes also included. The genes without numbers or coloured grey were not found to be syntenic across genera. From the left: 1 – catabolic 3-dehydroquinase; 2 – 3-dehydroshikimate dehydratase; 3 – quinate dehydrogenase; 4 – quinate permease; 5 – hypothetical protein; 6 – *CDO4*; 7 – Hypothetical protein part of fungal TF MHR superfamily, containing a GAL4 zinc finger DNA-binding domain; 8 – phosphopantetheine adenylyltransferase; 9 – subtilase family protein; 10 – DNA polymerase kappa; 11 – hypothetical protein part of beta elim lyase superfamily; 12 – subtilase/peptidase family; 13 – subtilase/peptidase family ; 14 – histone-lysine N-methyltransferase SET9; 15 – FK506-binding protein 2.

### Analysis of *CDO* gene family evolution

Parsimony analysis of the *CDO*s showed a simple evolutionary history of the genes (Figure 6). Based on the results of the parsimony analysis, a loss of the *CDO1* gene occurred in the common ancestor of *Bretziella, Ambrosiella* and *Huntiella.* This gene was also independently lost in the genus *Chalaropsis* when it split from the *Ceratocystis. Huntiella* incurred a simultaneous loss of *CDO2* and *CDO4* after it split from its closest relatives, the genera *Ambrosiella* and *Bretziella*. *CDO2* was also lost in *Ambrosiella* after diverging from *Bretziella*. A duplication of *CDO4* occurred in the common ancestor of *E. polonica* and *E. laricicola,* suggesting that they are paralogs which have continued to diverge over time (Figure 6).

**Figure 6:**
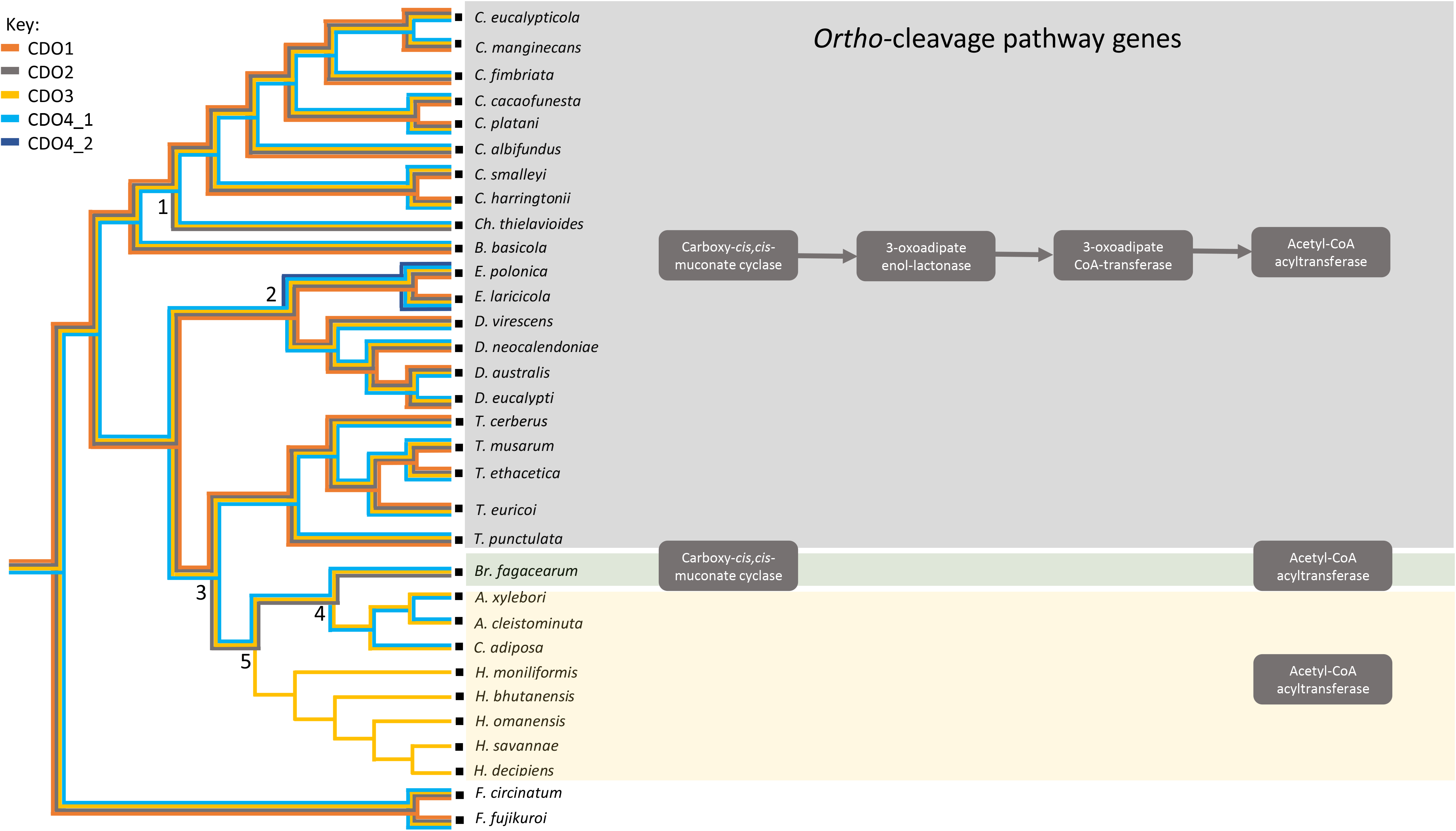
Ancestral history of the *CDO 1-4* genes (left image) based on a multi-locus species tree. The presence of a gene is indicated by the coloured line (*CDO1* – orange line, *CDO2* – grey line, *CDO3* – yellow line, *CDO4* – light blue line, *CDO4* duplication – dark blue line), and the loss of a gene is indicated by the absence of that coloured line from the node. Numbers on graph represent gene loss/gain events (1 – loss of *CDO1;* 2 – gain of *CD04*; 3 – loss of *CDO1*; 4 – loss of *CDO2*; 5 – loss of *CDO4*). Graphical representation (right image) of *ortho*-cleavage pathway genes present in each species.

An alternative evolutionary history was determined using a Count analysis, which determines the distribution of homolog family sizes across the different genera using a Wagner parsimony approach (Figure S2). The predicted gains and losses on the tree (Figure S2) showed that many were lineage specific. Consistent with the previous results, the ancestral state of the genes predicted that the most recent common ancestor of the Ceratocystidaceae contained a single copy of each of the CDOs. A single gene loss for *CDO1* was predicted for the *Ch. thielavioides* (Figure S2). A gene loss event of both *CDO1* and *CDO2* was predicted to have occurred in the lineage leading to the *Huntiella, Bretziella* and *Ambrosiella* lineages, which included *Ca. adiposa*. In contrast to the ancestral state reconstruction in Mesquite using parsimony, a gene gain of *CDO2* was predicted to have occurred for *Br. fagacearum.* The gene expansion predicted for the *Endoconidiophora* species was consistent with that of the method used in Mesquite.

When observed as a whole, the gene loss events were found to have occurred in the clades containing the saprophytic fungi and weakly pathogenic fungi, while the pathogenic fungi contained all four *CDO* copies (Table 1 and Figure 6). The Fisher’s Exact Test showed an association between *CDO* copy number and lifestyle of the Ceratocystidaceae (p < 0,01).

### Identification of the *ortho-* and *meta-*cleavage pathways in the Ceratocystidaceae

The genes for the entire *ortho-*cleavage pathway in the Ceratocystidaceae were identified in all of the genera except for those of *Huntiella, Ambrosiella, Catunica* and *Bretziella* (Table 2). There was one exception, the acetyl-CoA acyltransferase gene (EC 2.3.1.16) was present in all Ceratocystidaceae represented genomes (Table 2). The genes involved in the entire *meta-*cleavage pathway could not be identified in any of the fungal species, apart from one gene, a putative aldehyde dehydrogenase (EC 1.2.1.85).

**Table 2:** Identification of ortho-cleavage genes involved in the degradation of catecholic substrates in the genomes of the Ceratocystidaceae

| *Ortho*-cleavage pathway genes |  |  |  |  |
| --- | --- | --- | --- | --- |
| Species | Carboxy-<br>*cis,cis*-<br>muconate<br>cyclase | 3-oxoadipate<br>enol-lactonase | 3-oxoadipate<br>CoA-transferase | Acetyl-CoA<br>acyltransferase |
| *Ambrosiella cleistominuta* |  |  |  | √ |
| *A. xylebori* |  |  |  | √ |
| *Berkeleyomyces basicola* | √ | √ | √ | √ |
| *Bretziella fagacearum* | √ |  |  | √ |
| *Catunica adiposa* |  |  |  | √ |
| *Ceratocystis albifundus* | √ | √ | √ | √ |
| *Ce. eucalypticola* | √ | √ | √ | √ |
| *Ce. cacaofunesta* | √ | √ | √ | √ |
| *Ce. fimbriata* | √ | √ | √ | √ |
| *Ce. harringtonii* | √ | √ | √ | √ |
| *Ce. manginecans* | √ | √ | √ | √ |
| *Ce. platani* | √ | √ | √ | √ |
| *Ce. smalleyi* | √ | √ | √ | √ |
| *Chalaropsis thielavioides* | √ | √ | √ | √ |
| *Davidsoniella australis* | √ | √ | √ | √ |
| *D. eucalypti* | √ | √ | √ | √ |
| *D. neocaledoniae* | √ | √ | √ | √ |
| *D. virescens* | √ | √ | √ | √ |
| *Endoconidiophora laricicola* | √ | √ | √ | √ |
| *E. polonica* | √ | √ | √ | √ |
| *Huntiella bhutanensis* |  |  |  | √ |
| *H. decipiens* |  |  |  | ✓ |
| *H. moniliformis* |  |  |  | ✓ |
| *H. omanensis* |  |  |  | ✓ |
| *H. savannae* |  |  |  | ✓ |
| *Thielaviopsis cerberus* | ✓ | ✓ | ✓ | ✓ |
| *T. ethacetica* | ✓ | ✓ | ✓ | ✓ |
| *T. euricoi* | ✓ | ✓ | ✓ | ✓ |
| *T. musarum* | ✓ | ✓ | ✓ | ✓ |
| *T. punctulata* | ✓ | ✓ | ✓ | ✓ |

\*Reference genes used in BLAST query: Carboxy-*cis,cis*-muconate cyclase (*Ce. fimbriata* PHH5432.1); 3-oxoadipate enol-lactonase (*Ce. platani* KKF95260.1); 3-oxoadipate CoA-transferase (*Ce. platani* KKF92536.1); acetyl-CoA acyltransferase (*Ce. fimbriata* PHH52801.1).

## Discussion

Throughout the fungal kingdom, closely related species often differ significantly in their lifestyles (Soanes et al. 2008). This is also observed in the Ceratocystidaceae, a family which includes fungi that are associated with plants in different ways, ranging from saprophytic to highly pathogenic. Different lifestyles in closely related fungi are often determined by different gene inventories and not by evolutionary distances (Schäfer et al. 1989; Yoder and Turgeon 2001).

Plants synthesize a wide range of phenolic compounds, usually in high concentrations (Levin 1971). These compounds have antifungal activity, through binding to and precipitating extracellular proteins (Charlton et al. 2002; Ndhlala et al. 2015), interfering with membrane potentials (Rao et al. 2010), disrupting membrane structure (Sang Sung and Gun Lee 2010) and the oxidative respiratory electron chain (Shalaby and Horwitz 2015). In order to survive in association with plants, as epiphytes, pathogens or endophytes, fungi have adapted to these compounds, and intricate mechanisms to circumvent their toxicity have evolved (Anderson et al. 2010; Frantzeskakis et al. 2019).

An adaptation to phenolic plant defences are catabolic degradation pathways, where the phenolic ring is cleaved by a catechol dioxygenase enzyme and the linearized carbon chain is subsequently used as an energy source (Broderick 1999). In previous studies, CDO enzymes were identified as pathogenicity and virulence factors in the bark-beetle associated blue-stain fungus, *E. polonica,* a member of the Ceratocystidaceae (Hammerbacher et al. 2013; Wadke et al. 2016). To expand on previous research, CDO genes from different genera of the Ceratocystidaceae with widely different ecological lifestyles were investigated in this study with a focus on genomic differences using a comparative approach.

### *CDO* genes in the Ceratocystidaceae are more abundant in pathogenic species

In previous studies, four CDOs were described in *E. polonica,* (CDO1-CDO4) (Wadke et al. 2016). In this study, all four homologous CDOs were identified in the genera *Ceratocystis, Davidsoniella, Berkeleyomyces, Endoconidiophora* and *Thielaviopsis*. Species in these genera are considered highly pathogenic on diverse hosts. For example, *Ce. fimbriata* causes black rot of sweet potatoes, *Ce. manginecans* and *Ce. albifundus* cause wilt in Acacia trees and *Ce. platani* causes canker stains in plane trees (Halsted and Fairchild 1891; Johnson et al. 2005; Ocasio-morales et al. 2007; Roux et al. 2007; Tarigan et al. 2011; Adawi et al. 2013). *Thielaviopsis* species cause disease on many species of palm trees around the world (Paulin-Mahady et al. 2002; Melo et al. 2016; Saeed et al. 2016) and *Be. basicola* causes root rot in many hosts such as cotton, chicory, carrots, groundnuts and tobacco plants (Punja and Sun 1999; Nel et al. 2017). *Endoconidiophora* species are vectored by bark beetles to coniferous hosts such as *Picea abies* and *Larix decidua* causing blue-stain of the bark and wood, and kill the trees under high inoculation densities (Krokene and Solheim 1997; Krokene and Solheim 1998). The genus *Davidsoniella* mostly contains pathogenic species. *Davidsoniella virescens* is a virulent pathogen, causing sap streak in maple trees, but is also weakly pathogenic on other hardwood hosts (Shigo 1962; Richter 2012; Bal et al. 2013). In contrast, *D. eucalypti* is weakly pathogenic to certain species of *Eucalyptus* (Kile et al. 1996; Richter 2012; Wingfield et al. 2018), however, as it was only isolated from artificial stem wounds its primary host is not known.

In contrast to the pathogenic genera included in this study, only one CDO was identified in our study from the genus *Huntiella*. Species of this genus are saprophytic and colonise fresh wounds in tree bark, without causing disease (Roux et al. 2004; Van Wyk et al. 2006). This dramatic gene loss in the genus *Huntiella* supports previous findings, that the degradation of phenolic compounds is more common in pathogenic fungi compared to their saprophytic counterparts (Gluck-Thaler and Slot 2018). This pattern of gene loss in the saprophytic species of the Ceratocystidaceae has also been observed in the genes of the glycoside hydrolase protein family, which are involved in the catabolism of sucrose (Van der Nest et al. 2015).

*Ambrosiella* species, which are the mycangial symbionts of ambrosia beetles (von Arx and Hennebert 1965; Mayers et al. 2015; Mayers et al. 2019) only retained two of the four *CDOs* found in the Ceratocystidaceae. As ambrosia beetle symbionts, *Ambrosiella* species are also only weakly pathogenic, but are required to assimilate sufficient nutrients from the host to sustain the beetles’ development (Hulcr and Dunn 2011; Jankowiak 2011). *CDO1* and *CDO2*, therefore, do not seem to play an important role in nutrient acquisition for these fungi. Furthermore, the genome of *Ca. adiposa*, formerly *Ceratocystis adiposa* (Mayers et al. 2019), also contained only two *CDOs* (*CDO3* and *CDO4*) and is an opportunistic pathogen, which causes root rot in sugarcane cuttings but is considered a weak pathogen and does not attack established plants (Sartoris 1927). Furthermore, *CDO1* was lost from the genome of *Ch. thielavioides*. This species has a similar lifestyle to *Ca. adiposa* and is an opportunistic pathogen causing black mould in weakened plants, such as rose grafts (Longree 1940).

*CDO1* was also lost from the genome of *Br. Fagacearum,* a highly virulent pathogen, causing red oak wilt (Juzwik et al. 2008; de Beer et al. 2017). However, this fungus, which enters the tree through wounds and root grafts, restricts itself to the xylem vessels of the outer layers of the sapwood and avoids contact with phenolic-producing parenchyma cells (Juzwik and French 1983; Appel et al. 1987). Therefore, CDO copy number and type in the Ceratocystidaceae is not only determined by pathogenicity, but also by the pathogen’s mode of infection, its preferred niche within the host and the resilience of the host tissue.

Although there does appear to be a trend in *CDO* copy number and ecological lifestyle, it is clear that the gene loss is largely restricted to a single clade with fewer pathogens (Figure 6). As such the variation of the distribution of *CDOs* observed could be due to phylogenetic relatedness. To further investigate this, future studies will need to include more species, providing multiple independent pathogen and non-pathogen clades that could be compared for *CDO* copy number distribution.

### The loss of the *ortho-*cleavage pathway in the Ceratocystidaceae

Another contributing factor to the differences in mode of infection among the Ceratocystidaceae representatives may be attributed to the complete absence of the *ortho-*cleavage pathway in the genera that have lost some of their intradiol CDOs. The species which have lost this pathway are all grouped in one clade suggesting that the loss of this pathway is in some way related to their evolution over time. The clade includes the species of *Huntiella, Bretziella, Ambrosiella* and *Catunica*.

The absence of the *ortho-*cleavage pathway in *Huntiella, Catunica, Ambrosiella* and *Bretziella* will require further research to understand if the gene loss in these genera provides a fitness disadvantage for these fungi and limits them to their specific ecological niches. The *ortho-*cleavage pathway is predominantly used by fungi to degrade phenolic compounds (Leatham et al. 1983; Camarero et al. 1994; Fountoulakis et al. 2002; Shanmugam et al. 2010). As the *ortho-*cleavage pathway produces intermediates which enter the β-ketoadipate pathway, it also provides a nutritional advantage to the fungus (Hammerbacher et al. 2013; Wadke et al. 2016).

The single *CDO* present in *Huntiella* provides interesting insight into some differences between the pathogenic and saprophytic species. The lack of synteny observed between the flanking regions surrounding *CDO3* of *Huntiella* and other genera of the Ceratocystidaceae provides further evidence of differences in the evolutionary histories of the gene in the different species. Discordance in local synteny has been found to indicate non-orthologous gene relationships (Jun et al. 2009) and may contribute to the lifestyle adaptations of fungi (Yoder and Turgeon 2001). A lack of local synteny surrounding genes involved in pathogenicity in the Ceratocystidaceae has also been observed for genes involved in the utilization of plant-derived sucrose (Van der Nest et al. 2015), where the gene loss and lack of synteny were attributed to the action of transposable elements (TEs) found in the Ceratocystidaceae genomes. TEs have been shown to have an important impact on fungal lifestyles (Grandaubert et al. 2014; Van der Nest et al. 2015; Muszewska et al. 2019).

Although the *meta-*cleavage pathway has not been characterized in fungi, the presence of the extradiol dioxygenase (*CDO3*) in the *Huntiella* genomes and the increase in expression of CDO3 in *H. moniliformis* when grown in the presence of caffeic acid indicates that it does utilise the enzyme (data not shown). In *E. polonica* expression of CDO3, which was shown to have ring-cleaving activity *in vitro*, also increased during spruce infection (Wadke et al. 2016). As the complete *meta-*degradation pathway has not been resolved in fungi, there are two possible scenarios. The first is that all the fungi in this study can fully metabolise the phenolics using an as yet uncharacterised pathway involving extradiol cleavage. The second scenario is that the extradiol enzyme detoxifies the plant secondary metabolites through cleavage, promoting the survival of the fungus, but does not fully metabolise them (Yu and Keller 2005). Such a scenario was observed in the saprophyte *Ophiostoma piceae* which was able to initially degrade, but not metabolise, monoterpenoids to ensure its continued survival (Yu and Keller 2005; Haridas et al. 2013). Further investigation into the importance of extradiol dioxygenases in fungal plant infection should be undertaken to determine how they are utilised, as well as their role during plant infection. Due to their presence across all the species of the Ceratocystidaceae, it is likely that the extradiol dioxygenases are important for the metabolism of both the pathogenic and saprophytic species.

## Conclusions

In this study we considered the evolution of *CDOs* in a subset of species of the Ceratocystidaceae and their potential links with fungal lifestyle. Publicly available genomes were used to identify the full complement of genes encoding catechol dioxygenase enzymes in multiple species of the Ceratocystidaceae. While the genomes of the necrotrophic pathogens, with the exception of *Be. fagacearum,* contained four different genes encoding *CDOs*, mildly pathogenic species only contained two to three genes and saprophytic species only contained a single gene. The loss of the genes and the associated metabolic pathway appears to have occurred in a lineage specific manner.

Further investigation into the ecological importance of the CDOs in a wider range of species from the Ceratocystidaceae needs to be conducted to determine which CDOs are favoured by species that occupy different ecological niches. Future studies should also include the analysis of *CDO* gene expression while the fungi are occupying their specific niches and enzyme activity assays. This information will further aid in understanding the importance of CDOs in plant pathogen fitness and evolution.

## Abbreviations

CDO: Catechol dioxygenase
TCA: Tricarboxylic acid cycle

## Declarations

### Ethics approval and consent to participate

Not applicable.

### Availability of data and materials

The datasets used and/or analysed in the current study are available in the NCBI Database (https://www.ncbi.nlm.nih.gov/nucleotide/).

## Acknowledgements

We would like to acknowledge University of Pretoria and the Forestry and Agricultural Biotechnology Institute for use of infrastructure and the National Research Foundation for funds to BDW (DSI/NRF SARChI Chair for Fungal Genetics) and the Max Planck Society for funds to AH.

